# A ventral tegmental area GABAergic projection to the ventral pallidum regulates value-based decision making in mice

**DOI:** 10.64898/2026.02.19.706858

**Authors:** Wen-Liang Zhou, Hanna Yousuf, Yann S. Mineur, Marina R. Picciotto

## Abstract

Activity of the mesolimbic system is essential for adaptive performance of reward-related behaviors. Within this system, dopaminergic (DAergic) neurons play a critical role in driving motivation to obtain rewards and encoding predictions and error signals during reinforcement learning. However, activity of DAergic neurons shifts from reward presentation to predictive cues following cue-reward learning, leaving open questions about the mechanism of subjective reward value representation. Our previous studies suggest that activity of a GABAergic circuit originating from the ventral tegmental area (VTA) and projecting to the ventral pallidum (VP) scales with unconditioned reward value, independent of effort or associative cue-reward learning. Here, we demonstrate that activity in this pathway consistently reflects unconditioned reward value across extended cue-reward training, unlike DA activity, which undergoes dynamic changes toward the cue and away from a predicted reward. VTA-to-VP GABA activity tracks internal-state-dependent reward value, showing minimal response to water drinking in sated mice and strong activity after overnight dehydration. In a two-option probabilistic operant reward task (PRT), optogenetic activation of this pathway upon reward consumption biased decision-making toward the stimulation-paired option, even when its reward was of lesser value. These findings identify a previously uncharacterized circuit that encodes reward value and contributes to value-based decision-making.

**Significance Statement:** Adaptive behavior depends on accurate representation of reward value. While dopamine (DA) signals shift from rewards to predictive cues during learning, the neural encoding of unconditioned reward value has remained elusive. We identify a GABAergic projection from the VTA to the VP that stably encodes unconditioned reward value across extended training, yet tracks changes in internal state such as thirst. Unlike DA activity, this pathway consistently reflects reward consumption and, when stimulated, biases choice toward otherwise less-preferred options. These findings uncover a stable but state-sensitive mechanism of reward value encoding, providing a new framework for understanding value-based decision-making and its disruption in neuropsychiatric disorders.

## INTRODUCTION

Survival fundamentally depends on the value an individual derives from rewards earned through adaptive behaviors (Schultz 2015, Juechems and Summerfield 2019). To guide these actions, the brain must encode the current value of a reward. Optimal behavioral outcomes are believed to occur when choices are made based on the option with the highest value in a given reward-related context (Tajima, Drugowitsch et al. 2016).

Importantly, reward valuation has been linked to cognitive processing (Kobayashi and Hsu 2019, Loganathan 2021), and is affected in a number of psychiatric disorders, including depression, schizophrenia, and drug addiction (Kalivas and Volkow 2005, Rupprechter, Stankevicius et al. 2021). Given the fundamental importance of encoding outcome value for survival, mammalian brains have evolved complex neural mechanisms to compute and represent different types of value. For instance, activity in the orbitofrontal cortex represents the value of reward-predictive cues (Kahnt, Heinzle et al. 2010); the nucleus accumbens (NAc) signals value predictions as a function of dopamine-encoded prediction error signals in the ventral tegmental area (VTA) (Mannella, Gurney et al. 2013); and the parietal cortex encodes the value of future actions (Sugrue, Corrado et al. 2004). However, it remains unclear how the unconditioned, “intrinsic” value of a reward is represented in the brain, and how this encoding contributes to reward-related behaviors such as value-based decision-making. Furthermore, it is possible that the encoding of different types of value—including unconditioned reward value, economic values (e.g., probability, effort, delay), and action value—may converge within a common circuit to generate a final subjective (current) value that shapes learning and choice.

Classically, midbrain DAergic neurons have been proposed to encode reward prediction errors (Bayer and Glimcher 2005, Schultz 2015), such that phasic responses to reward outcomes diminish to baseline as rewards become fully predicted through learning.

Consistent with this framework, DA responses to primary rewards often decrease with training as predictive cues acquire informational value. However, recent work has revealed substantial heterogeneity among DAergic neurons (Lammel, Lim et al. 2014, Azcorra 2023). A subset of DAergic neurons continues to respond robustly to predicted rewards, and in some behavioral tasks, reward-evoked dopamine signals do not progressively decline but instead reach a stable, plateau-like level with training (Kim 2024). These findings suggest that dopamine signaling cannot be fully captured by a uniform RPE model. Nevertheless, the prevailing view is that dopamine reward responses are highly variable across neurons, task structures, and motivational states, and therefore may lack the stability required to reliably encode unconditioned reward value (Lerner, Holloway et al. 2021, Blanco-Pozo, Akam et al. 2024). Rather than representing a fixed intrinsic value, dopamine activity appears to be dynamically shaped by learning and context, limiting its suitability as a stable reference signal for reward valuation.

We previously showed that activity of a GABAergic pathway originating in the VTA and projecting to the ventral pallidum (VP) correlates consistently with the value of a reward during consumption (Zhou, Kim et al. 2022), consistent with emerging evidence that VTA GABAergic neurons, at both population and projection-specific levels, encode reward-related and motivational signals (Yoo 2016, Bouarab 2019, Stelzner 2025). Optogenetic activation of this pathway enhanced performance in a cued reward task, suggesting that this circuit may encode the unconditioned value of a reward as it is consumed. In the current study, we therefore investigated whether VTA-to-VP GABA responses remain stable across time and experiences that alter reward salience. Such stability would provide a reliable internal signal for evaluating rewards relevant to survival.

Stable value representation is also critical for computing prediction errors during associative learning, which requires a reference signal that is not itself altered by the learning process (Watabe-Uchida, Eshel et al. 2017). Importantly, the value of a reward depends on how effectively it supports survival and reproduction and can vary dramatically with physiological state (Minamimoto, Yamada et al. 2012, Juechems and Summerfield 2019). To test this, we measured VTA-to-VP GABA activity during water consumption in both satiated and dehydrated states. Finally, we employed precisely timed optogenetic stimulation during a two-option probabilistic reward task (PRT) to determine whether manipulating value-encoding activity could bias choice preference, providing causal evidence that this pathway encodes reward value and can override natural reward valuation.

## RESULTS

### Consistent VTA-to-VP GABA responses to reward consumption across long-term Pavlovian Conditioning, compared to DA transmission in the NAc shell

Midbrain DAergic neurons are known to signal predictions and prediction errors during associative reward learning, with neuronal response to the same rewarded outcome shifting over the course of learning toward a randomly-delivered, reward-predictive cue (Schultz 2015). In rats, DA transmission in the NAc core aligns with prediction-error signaling (Saddoris, Cacciapaglia et al. 2015), whereas in the NAc shell DA tracks the value of the rewarded outcome (Sackett, Saddoris et al. 2017). We therefore measured the dynamics of DA transmission in the NAc shell during associative learning in mice using fiber photometry. Briefly, a genetically encoded fluorescent DA sensor (AAV-hsyn-DA4.4) (Hamilos, Spedicato et al. 2021) was expressed in the NAc shell of GAD65-Cre mice, and an optic fiber was implanted targeting the site of viral infusion. After recovery, lightly restrained mice underwent Pavlovian conditioning for 25 sessions (5 sessions per week, Fig. S1). The light restraint paradigm ensured consistent timing of cue presentation and reward consumption across trials, enabling reliable alignment of neural activity to US onset with minimally-distorted averaging of CS-related responses (Fig. 1A). The conditioned stimulus (CS) was presented on a random time 60-s (RT-60) schedule, and 5 s following CS onset, a 2-s liquid reward (Ensure®) was delivered as the unconditioned stimulus (US).

**Figure 1.**
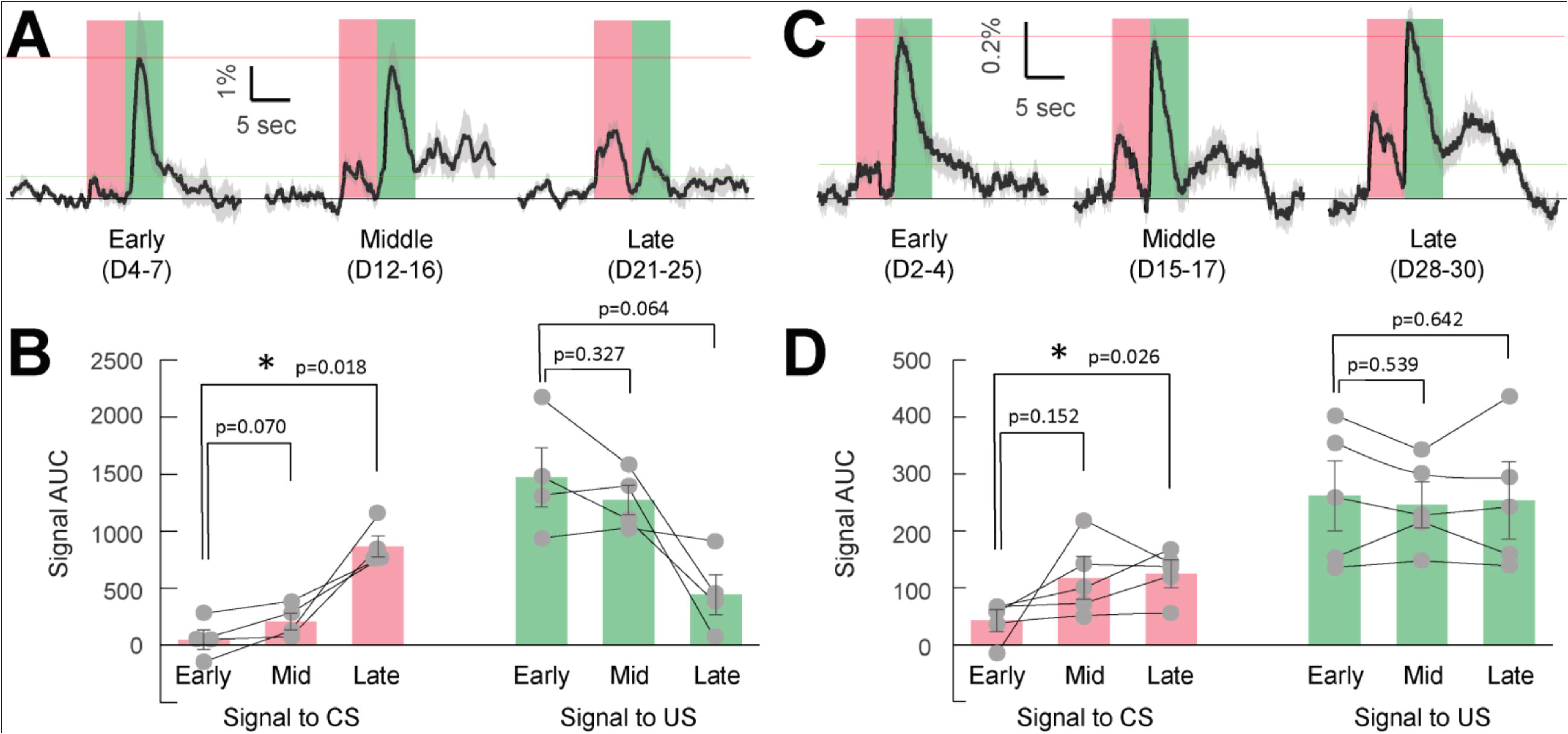
Dynamics of DA transmission in the NAc shell vs. VTA-to-VP GABA activity during Pavlovian conditioning. (A) DA sensor DA4.4 was expressed in the NAc shell to monitor fluorescence changes (dF/F), serving as an indicator of DA transmission during Pavlovian conditioning at Early (Day 4-7), Middle (Day 12-16), and Late (Day 21-25) stages of training. Fluorescence signals were aligned to the onset of US delivery, indicated by the left edge of the green shaded area. The onset of the CS is indicated by the left edge of the red shaded area. The red portion of the signal trace (-5 to 0 s) was used to calculate the area under the curve (AUC) for CS-evoked DA responses, while the green portion (0 to 5 s) was used to compute the AUC for US-evoked DA responses. Signals were averaged from 4 male mice from the indicated training sessions. (B) Quantification of signal AUCs from (A). The left red bars show CS-evoked responses, and the right green bars show US-evoked responses across Early, Middle, and Late training stages. Each dot represents the averaged signal AUC measured in one mouse in each stage. Statistical comparisons between conditioning stages were conducted using Student’s t-test. N = 4 male mice. Based on the Early and Late signals to CS in this dataset, a paired t-test power analysis (α = 0.05, β = 0.20) indicated that a minimum of 3 animals would be sufficient. (C) GCaMP7s was expressed in the VTA GAD65+ neurons and GCaMP fluorescence was recorded from the axon terminals projected into the VP, during Pavlovian conditioning at the Early (Day 2-4), Middle (Day 15-17), and Late (Day 28-30) stages. Signal analysis was performed the same as described in (A). (D) Quantification of signal AUCs from (C). The left red bars show CS-evoked responses, and the right green bars show US-evoked responses across training stages. Student’s t-test was used to assess statistical significance among conditioning stages. N = 5 male mice. Based on the Early and Late signals to CS in this dataset, a paired t-test power analysis (α = 0.05, β = 0.20) indicated that a minimum of 5 animals would be sufficient.

During the early phase of training (Day 4-7), we observed robust, time-locked DA signals in response to US delivery/consumption (Fig. 1A,B; signal AUC: 1472 ± 258; n = 4 mice). As training progressed, US-related DA signals declined during the late phase (AUC: 442 ± 177; D21-25), although this did not reach statistical significance (Student’s t-test, p = 0.064). In sharp contrast, DA signals in response to CS increased significantly (Early: 48 ± 81; Late: 845 ± 92; p = 0.018). The dynamics of DA transmission in the NAc shell are consistent with previously reported CS- and US-evoked DA responses following cue-reward training in primates (Schultz, Dayan et al. 1997).

We next measured VTA-to-VP GABA activity in this paradigm under the same conditions, except that mice were allowed to move freely in the box (Fig. S1C). We infused pGP-AAV1-syn-FLEX-jGCaMP7s-WPRE (Zhang and Looger 2024) into the VTA of GAD65-Cre mice and implanted an optical fiber above the VP (Fig. S3A-E). This enabled us to record calcium-dependent fluorescence signals from the terminals of VTA GAD+ neurons in the VP of behaving mice (Zhou, Kim et al. 2022). Similar to DA signals in the NAc shell, activity of VTA-to-VP GABA neurons was time-locked to reward consumption during early sessions (Fig. 1C&D; AUC:262 ± 61; D2-4; n = 5 mice). As training progressed, GABA activity remained stable across 30 training sessions, showing negligible changes in amplitude or duration (Late phase AUC: 254 ± 68; D28-30; p = 0.64). We also observed that the GABA response to the CS (transients red shaded, Fig. 1C&D) increased from the Early phase (AUC: 43 ± 19) to Late phase (AUC: 125 ± 24; p = 0.026), largely mirroring the cue-dependent dynamics of DA activity. These results suggest that VTA-to-VP GABA activity encodes the stable value of unconditioned rewards, while also reflecting the acquired value of predictive cues following conditioning.

As expected, training resulted in effective Pavlovian conditioning in both groups of mice. Following training, both groups showed a significant preference for the reward-paired cue in conditioned reinforcement tests, indicating robust and comparable acquisition of cue–reward associations across conditions (Fig. S2).

### VTA-to-VP GABA neurons track the intrinsic value of water under conditions of satiation and dehydration

The value of a reward depends on the extent to which it aids reproduction and survival, and can vary greatly depending on physiological condition or internal state (Schultz 2015, Juechems and Summerfield 2019). For instance, thirst triggers a motivational drive and therefore increases the value of drinking water (Minamimoto, Yamada et al. 2012). Based on this idea, we determined whether VTA-to-VP GABA activity might reflect differences in the value of drinking water under satiated and dehydrated conditions. We expressed GCaMP7s in the VTA of GAD65-Cre mice and implanted an optical fiber above the VP to monitor calcium activity from GABAergic axon terminals. Following recovery, freely-moving mice underwent Pavlovian conditioning to earn rewards (Ensure) or drinking water from a receptacle.

On Day 1, Ensure consumption elicited a robust, time-locked calcium response in VTA-to-VP GABA terminals (AUC: 379 ± 48; N = 5 mice; Fig. 2A&D). On Day 2, the same volume of water was delivered in place of Ensure. In mice that had not experienced water deprivation, calcium signals during water licking were negligible (AUC: -44 ± 50; Fig. 2B&D). Mice were then deprived of water for 18 h until the next testing session. On Day 3, water consumption evoked significantly greater calcium responses in VTA-to-VP GABA terminals compared to the water-sated condition (AUC: 267 ± 37; p = 0.020; Fig. 2C&D), although responses remained lower than those elicited by Ensure (p = 0.011). These results indicate that VTA-to-VP GABA activity tracks the intrinsic value of water, which depends on the animal’s internal physiological state.

**Figure 2.**
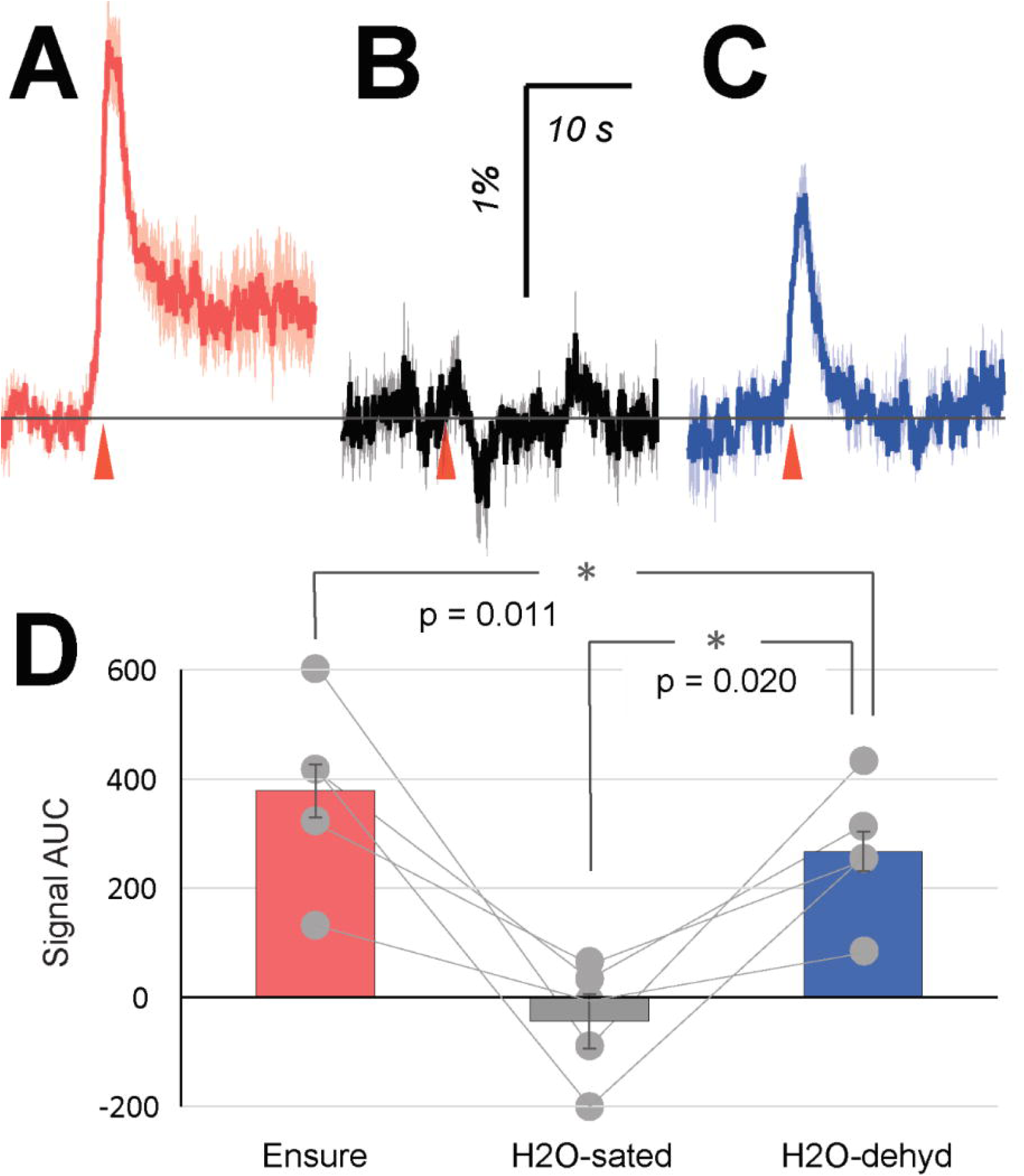
VTA-to-VP GABA activity in response to water under satiation and dehydration. GCaMP7s was expressed in VTA GAD65+ neurons, and GCaMP fluorescence was recorded from the axon terminals projected into the VP during Pavlovian conditioning. (A) On Day 1, Ensure was delivered into the receptacle as a reward. GCaMP signals were averaged by aligning traces to the first receptacle entry (red arrowhead) following each reward delivery. (B) On Day 2, water was delivered into the receptacle instead of Ensure. Prior to this session, the mice had been continuously provided with ad libitum access to water for at least 3 weeks. (C) Following the Day 2 session, the mice were deprived of water for 18 hours before the subsequent session on Day 3, during which water was again delivered as a reward. (D) Quantification of GCaMP signal AUC from panels (A-C). Each dot represents the average AUC from an individual mouse during the corresponding task session. Statistical significance between conditions was assessed using Student’s t-test. N = 5 male mice. Based on the signals measured under H_2_O-sated and dehydrated conditions in this dataset, a paired t-test power analysis (α = 0.05, β = 0.20) indicated that a minimum of 5 animals would be sufficient.

### A mouse model to assess value-based reward choice

Assessment of reward value is vital for survival. Thus, it is essential to determine and encode value accurately and to adjust behavior accordingly. If VTA-to-VP GABA activity contributes to representation of reward value, manipulating activity of these neurons should alter value-based behaviors such as decision-making. To test this hypothesis, we implemented a two-option probabilistic reward choice task (PRT) to determine how VTA-to-VP GABA activity influences choices between operant responses to obtain rewards of different value (Milienne-Petiot 2017).

Male GAD65-Cre mice and wild-type littermates were trained to nose-poke for Ensure delivered through a metal spout. This delivery design ensured that each drop of liquid was either consumed immediately or allowed to drip away, preventing contamination of the perceived value of subsequent rewards. Following pre-training to habituate the animals to the apparatus, two nose-poke apertures were positioned side by side (Fig. 3A), each corresponding to a distinct reward schedule. Option A resulted in a 75% chance of a high-value reward (Ensure, ∼24 μl delivered over 2 s) and a 25% chance of water, whereas Option B resulted in a 25% chance of Ensure and a 75% chance of water. Mice had *ad libitum* access to water before and throughout training, making water a low-value option in this task. Thus, Option A represented a higher-value reward delivery schedule, while Option B represented a lower-value schedule.

**Figure 3.**
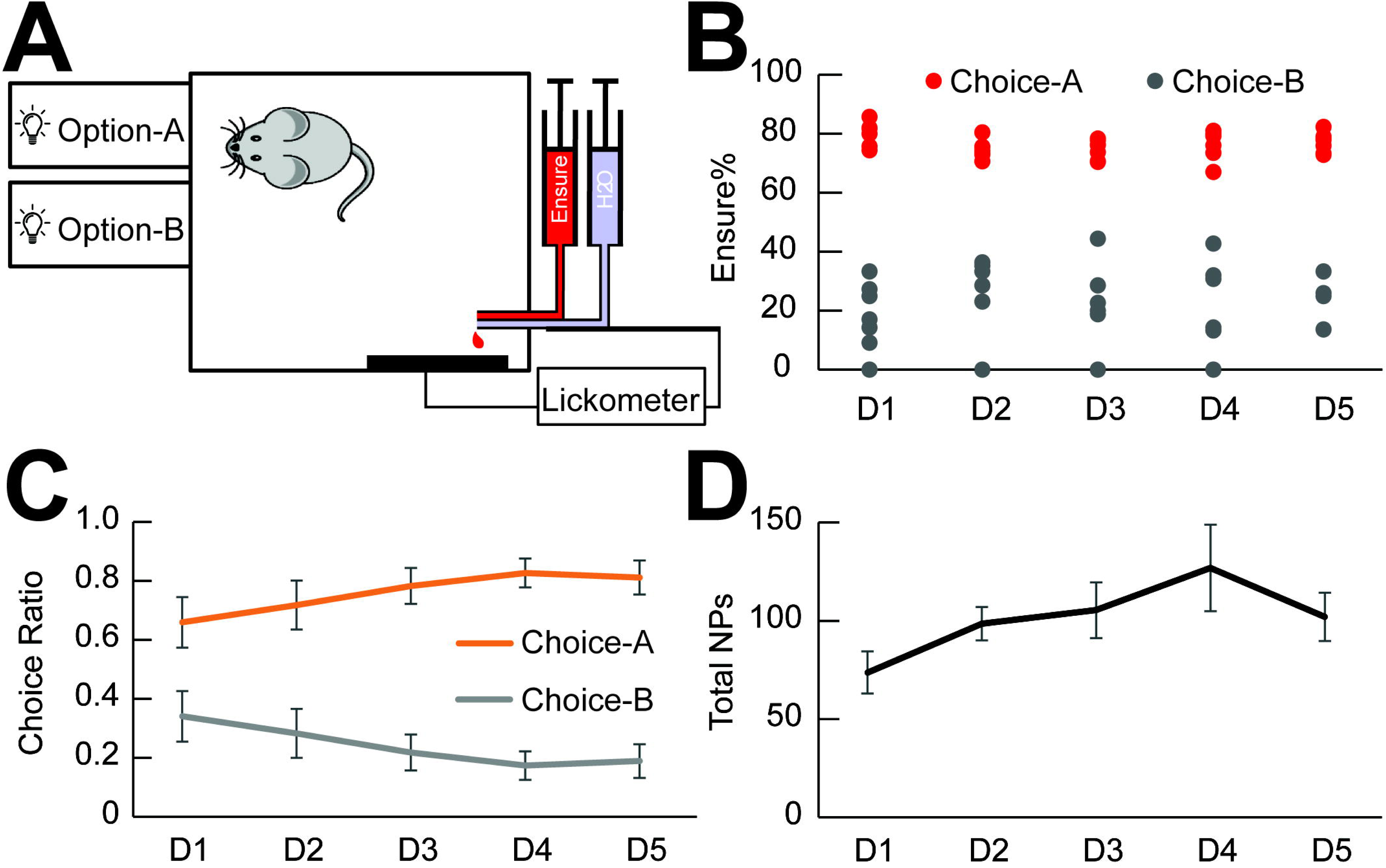
A mouse model of a two-option probabilistic reward choice task (PRT). Male GAD65-Cre mice (n = 5 mice) and their wild-type littermates (n = 2 mice) were trained in an operant task designed to assess value-based decision-making. (A) Schematic of behavioral chambers used in the PRT. In Option A, a nose-poke made into aperture A resulted in a 75% chance of receiving a high-value reward (Ensure, ∼24 μl, delivered over 2 s) and a 25% chance of receiving the same amount of water. Both liquids were delivered via a composite spout located at the lower-right corner. In Option B, a nose-poke into aperture B led to a 25% chance of receiving Ensure and a 75% chance of receiving water. The timing of each licking on the spout was recorded with a lickometer. (B) Actual recorded probabilities of receiving Ensure following Choice A (set at 75%) and Choice B (set at 25%), confirming task implementation. (C) Choice behavior across 5 training sessions (Days 1-5). Mice increasingly favored Choice A. Two-way ANOVA with repeated measures shows a significant main effect of choice (F(1,12) = 33.7, P = 8.4 × 10^-5^), and a significant effect of training session (F(4,24) = 5.46, P = 0.0029). N = 7 mice. (D) Total number of nose-pokes across the five sessions. A trend toward increased nose-poking was observed, but the session effect did not reach statistical significance (F(4,24) = 2.41, P = 0.078).

During 5 consecutive sessions, Ensure or water were delivered reliably at the designated probabilities (Fig. 3B). Mice quickly distinguished between the two options and developed a strong preference for Option A (75% Ensure delivery) over Option B (25% Ensure delivery) from the first session (Fig. 3C, Two-way ANOVA with repeated measures, Overall effect of option: F(1,12) = 33.7, P = 8.4 × 10^-5^; n = 7 mice). This preference persisted and strengthened over the course of 5 sessions (Day effect: F(4,24) = 5.46, P = 0.0029). There was a trend toward an increase in the total number of effective nose-pokes over days, but this did not reach statistical significance (Fig. 3D, p = 0.078). These data suggest that the PRT paradigm used here effectively reflects ability to calculate differences in value of operant responses that result in differentially rewarded outcomes. Furthermore, mice adjusted their behavior based on reward value to achieve an optimal outcome, making this model suitable for assessing the function of reward-encoding neural circuits.

### Optogenetic stimulation of VTA-to-VP GABA pathway during reward consumption shifts choice preference

Based on previous findings that the activity of VTA-to-VP GABA neurons increases during reward consumption, we hypothesized that activity in this circuit may encode the unconditioned value of a reward. Consequently, manipulating activity of this GABAergic pathway might influence reward valuation, and thereby alter value-based behaviors such as decision-making. To test this hypothesis, we used an optogenetic approach to stimulate VTA-to-VP GABAergic axon terminals in the VP while animals performed the PRT paradigm. AAV2-EF1a-DIO-hChR2(H134R)-eYFP was infused into the VTA of GAD65-Cre male mice, and bilateral optical cannulae were implanted targeting the posterior VP (Fig. S4). Control mice received AAV2-EF1a-eYFP infusions into the VP and bilateral fiber implantation. After recovery, mice were trained to choose between two reward options in the PRT. To quantify choice preference, we calculated the ratio for Option B (25% Ensure/75% water) relative to the total number of effective nose-pokes, enabling us to track directional shifts toward Option B.

During the initial five sessions (D1–D5), behavioral boxes were configured as shown in Fig. 4A. The optic fiber was connected but no light stimulation was delivered, establishing baseline behavior. As expected, prior to stimulation mice displayed a strong preference for Option A over Option B (Fig. 4C,E), consistent with results in unmanipulated mice (Fig. 3C). On D6, nose-poke apertures were rearranged as shown in Fig. 4B, and mice were trained for one day to develop a mild preference for the location where the aperture was associated with the higher-value reward. From D7 to D11, the positions of the two apertures were swapped, and mice were trained on the PRT while receiving optical stimulation of VTA-to-VP GABA terminals during Option B trials. Specifically, across these 5 sessions, whenever a mouse chose Option B—regardless of whether Ensure or water was delivered—a 5-s train of blue light pulses (20 Hz, 465 nm) was triggered upon the first lick. ChR2-expressing mice showed a significant increase in their choice of Option B compared to baseline (Fig. 4C&E, Two-way ANOVA with repeated measures, F(1,14) = 26.7, P = 1.4 × 10^-4^, N = 8 mice).

**Figure 4.**
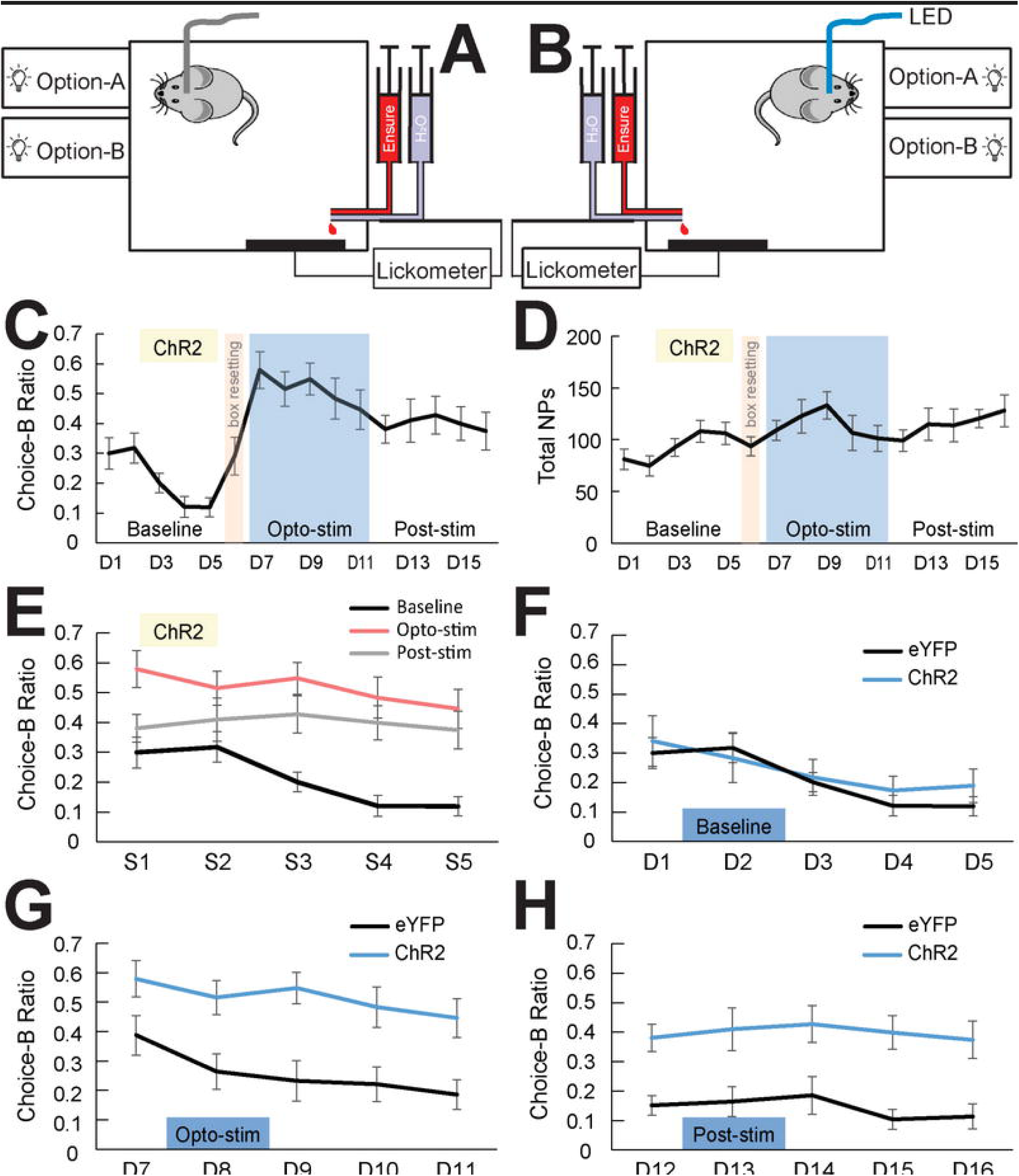
Stimulation of VTA-to-VP GABA pathway shifts choice preference in the PRT. (A) Schematic of the behavioral chamber used during baseline training in the PRT. (B) Setup used during the one-session transitional training of the PRT. Following this session, the spatial positions of Aperture-A and Aperture-B were reversed. This new configuration was used for all subsequent stimulation (Opto-Stim) and post-stimulation (Post-Stim) sessions. (C) Option-B choice ratio across Baseline, Opto-Stim, and Post-stim phases in mice expressing ChR2 in VTA GAD65+ neurons. The light brown shaded area represents the transitional session; the blue shaded area indicates five Opto-Stim sessions, during which blue light stimulation was delivered during reward consumption following Choice B. N = 8 mice. (D) Total number of effective nose-pokes recorded per session across the 3 task phases. (E) Option-B choice ratios across Baseline, Opto-Stim, and Post-stim phases in ChR2-expressing mice. Each phase includes sessions labeled as S1 through S5. Two-way ANOVA with repeated-measures revealed a significant increase in Option B choices during Opto-Stim compared to Baseline (F(1,14) = 26.7, P = 1.4 × 10^-4^), and a significant difference between Baseline and Post-Stim (F(1,14) = 8.32, P = 0.012). (F) Control group (n = 7 mice) expressing eYFP in VTA GAD65+ neurons, trained under the same PRT paradigm and schedule, as described in panel C. No significant difference was found between ChR2 and eYFP groups in Option B choice ratios during the Baseline phase (F(1,13) = 0.17, P = 0.68). (G) During the Opto-Stim phase, Option B choice ratios were significantly higher in ChR2 mice compared to eYFP controls (F(1,13) = 13.3, P = 0.0030). (H) Similarly, during the Post-Stim phase, ChR2 mice maintained a higher preference for Option B compared to controls (F(1,13) = 12.9, P = 0.0033).

Strikingly, this preference persisted after optical stimulation ceased, as mice continued to choose Option B more often than at baseline (Fig. 4C&E, F(1,14) = 8.32, P = 0.012). This persistent preference could reflect the formation of habitual or context-dependent reinforcement associated with the stimulation-paired option, rather than a sustained change in online reward-value computation.

As a control, GAD65-Cre male mice (N = 7) infused with AAV2-EF1a-eYFP into the VTA were tested in parallel with ChR2-expressing mice. During baseline sessions (D1-D5), both ChR2 and eYFP groups reliably discriminated between options of different values (Fig. 4F, F(1,13) = 0.17, P = 0.68). During Stimulation, ChR2-expressing mice exhibited a robust shift in preference toward Option B compared to eYFP controls (Fig. 4G, F(1,13) = 13.3, P = 0.0030). This stimulation-induced preference for Option B was sustained during post-stimulation sessions, in contrast to the stable preference for Option A observed in the eYFP group (Fig. 4H, F(1,13) = 12.9, P = 0.0033). Together, these results demonstrate that activity in the VTA-to-VP GABA pathway is sufficient to bias reward choice, overriding the natural preference for the higher-value option.

## DISCUSSION

Correctly calculating the value of a behavioral outcome is essential for survival and requires multiple neurobiological processes to reflect the magnitude of a delivered reward. Reward value depends on both external variables (e.g., utility) and internal motivational states (e.g., hunger or thirst), making it inherently subjective (Schultz 2015). A neural mechanism that encodes reward value should therefore integrate both external reward attributes and internal physiological states. Importantly, this subjectivity reflects systematic modulation by internal states, such that within a given motivational state, neural representations of reward value are expected to remain stable and reliable across time and experience (Enel 2020, Kimmel 2020, Ottenheimer 2023).

Organisms typically maintain a given physiological state for extended periods when seeking rewards such as food or water (Schultz 2015). Transitions between motivational states (e.g., from satiety to thirst) often occur in a nonlinear, threshold-like manner, following prolonged deprivation or consumption, and are characterized by relatively abrupt shifts in motivational drive (Becker 2017, Allen, Chen et al. 2019). Within each state, however, the intensity of the underlying need remains relatively stable (Keramati and Gutkin 2014, Encarnacion-Rivera 2025). Thus, an ideal value-encoding neural mechanism should be capable of consistent represention of the external attributes of the same reward, while the internal state remains constant. We refer to this stable, state-specific component of reward value as “unconditioned value”, assuming a relatively fixed internal physiological context. This concept is particularly important for cue-reward learning. According to reward prediction error theory, accurate encoding of the currently experienced reward is essential for the correct computation of prediction errors, which in turn guide adaptive behavior within a given motivational and environmental context (Schultz 2016, Nasser 2017).

We demonstrated previously that the activity of VTA-to-VP GABA projection neurons correlates with unconditioned reward value (Zhou, Kim et al. 2022). However, it remains unclear whether this pathway fully satisfies the criteria for encoding unconditioned value. Specifically, does it reliably reflect both the stable external variables and fluctuating internal states, and does its activity causally influence choice behavior? In the current study, we first used a classic Pavlovian conditioning paradigm to verify that the activity of VTA-to-VP GABA neurons remained stable and consistent over an extended period of cue-reward learning (30 training sessions over 40 days). This stable response pattern stands in contrast to the dynamic nature of DA signals in the NAc shell, which reflect the prediction of future reward and the error between expected and received value (Saddoris, Cacciapaglia et al. 2015, Schultz 2016, Lerner, Holloway et al. 2021). In the same cohort of mice, we measured DA signals in the NAc shell and observed a markedly different pattern from the VTA-to-VP GABA responses to identical rewards across multiple Pavlovian conditioning sessions. We also observed that VTA-to-VP GABA responses to the CS gradually increased over the course of training, resembling the evolving pattern of CS-responding DA signals. This suggests that VTA-to-VP GABA activity reflected the acquired value of the CS itself as the CS-US association formed during conditioning. Notably, the temporal dynamics of the CS-responsive signal in the DA and VTA-to-VP GABA pathways appeared to be qualitatively similar. Whether this similarity reflects functional interdependence between the two systems remains an open question and warrants further investigation.

Water is a prototypical reward whose value varies dramatically depending on hydration state. It is known that elevated blood osmolality levels increase motivation for water-seeking behavior (Minamimoto, Yamada et al. 2012), consistent with the idea that the subjective value of water increases substantially as an animal becomes dehydrated.

Using water as a reward, we showed that VTA-to-VP GABA activity tracked changes in water value across states of satiety and dehydration. In hydrated animals, VTA-to-VP GABA activity remained low, consistent with the minimal subjective value of water in sated mice. In contrast, when the same animals were water-deprived overnight, activity of the VTA-to-VP GABA pathway increased robustly, suggesting that VTA-to-VP GABA neurons effectively integrate both external variables and internal variables to encode the dynamic value of a reward.

In normative theories of value-based decision-making, an accurate representation of reward value serves as both the future criterion and the evidence used to guide decisions (Schultz 2015, Tajima, Drugowitsch et al. 2016). Optimal outcomes occur when choices are made based on the option with the highest value within a given reward-related context. Since mice can reliably perform value-based decision-making tasks (International Brain, Aguillon-Rodriguez et al. 2021), we designed a two-option probabilistic reward choice task (PRT) to determine whether the representation of reward value in VTA-to-VP GABA pathway might serve as a driving signal for decision commitment (Milienne-Petiot 2017). The probabilistic delivery of reward or water prevented the animals from becoming fixated on the higher-valued option, encouraging exploration of alternative choices. Mice from the Gad65-Cre line and their wild type littermates demonstrated a clear ability to discriminate between reward options of differing values, and rapidly developed a strong preference for the higher-valued Option A, even during the first session. This paradigm allows robust, quantifiable assessment of value-based decision-making. Leveraging this behaviorally grounded framework, we determined whether manipulating VTA-to-VP GABA activity could override appropriate reward valuation to alter value-based decision-making (Levy and Glimcher 2012)(Tom et al., 2019). Elucidating this mechanism could provide critical insights into how addictive substances, such as opioids—which lack intrinsic survival value—nonetheless exert a powerful influence on behavior by reshaping choice preferences.

According to normative theories of value-based decision-making, individuals compute the subjective value of each option and select the one with the highest expected value (Kording 2007, Summerfield and Parpart 2022). In the current study, stimulating the VTA-to-VP GABA pathway during reward consumption increased the value of the lower-rewarded Option B, shifting preference away from the higher-valued Option A. This supports the idea that the VTA-to-VP GABA pathway does not merely relay reward signals passively, but plays an active role in value attribution, which updates reward value during outcome evaluation, biasing future choices in line with value-updating models (Schultz 2015). In other words, stimulation of the VTA-to-VP GABA pathway alters the utility (value) function associated with reward Option B, thereby changing the expected value that guides future choice behavior (Kording 2007). The temporal specificity of the manipulation—restricted to the reward consumption phase—further supports the idea that post-decision valuation signals, rather than anticipatory or motor-related components, are critical for this modulation. This aligns with reinforcement learning models in which reward prediction errors and post-reward feedback are essential for updating internal value representations (Schultz 2015).

One potential alternative interpretation is that VTA-to-VP GABA neurons encode a more general motivational signal, such as incentive salience or stimulus intensity, rather than reward value per se. However, incentive salience is primarily associated with anticipatory, cue-driven “wanting” signals that operate before reward acquisition, directing attention and motivating approach behavior (Berridge 2018). In contrast, the manipulations and recordings in the current study were restricted to the reward consumption phase, when the animal was already engaged in consummatory behavior. At this stage, the dominant subjective experience is reward liking, reflecting the experienced hedonic value (liking) rather than anticipatory motivational salience (wanting) (Berridge and Robinson 2016, Berridge 2018). Moreover, salience-related signals are typically strongest in response to predictive cues and unexpected events (Schultz 2016, Berridge 2018), whereas effects reported here were observed during outcome evaluation, when the reward was already obtained and its sensory and hedonic properties were being processed. The ability of VTA-to-VP GABA stimulation during consumption to increase the subjective value of a lower-valued option selectively and bias future choice behavior therefore supports a role for this pathway in encoding reward value, rather than merely signaling salience or stimulus intensity.

Prediction-error calculation is one normative theory and is widely used to interpret cued-reward learning behaviors, especially during Pavlovian conditioning. In the mammalian brain, dopamine plays a central role in representing both the reward prediction for the CS and the surprise (or error) between the prediction and the rewarded outcome. These signals guide and constantly adjust behavior to optimize responses to reward-predicting cues. However, recent studies have shown that depletion of DA during cue-reward tasks does not disrupt learning outcomes (FitzGerald, Dolan et al. 2015), and stimulation of DA signaling does not alter future choice behavior (Blanco-Pozo, Akam et al. 2024). These results strongly suggest that DA signals are not sufficient to represent reward value as predicted by normative theories of value-based reward learning (Kording 2007, Summerfield and Parpart 2022). The current findings identify a novel neural mechanism that integrates both external and internal variables to encode reward value and actively modulates behavior for better (more highly rewarded) behavioral outcomes.

## Materials and Methods

### Animals

Male GAD65-IRES-Cre mice (GAD65-Cre; 4-10 months, The Jackson Laboratory) were group housed on a 12–12 h light–dark cycle with ad libitum access to food and water. All procedures were approved by the Yale University Institutional Animal Care and Use Committee and conducted in accordance with the NIH Guide for the Care and Use of Laboratory Animals. Age was matched and randomized across conditions within the same experiment.

### Surgery

For fiber photometry studies, GAD65-Cre mice were infused with pGP-AAV1-syn-FLEX-jGCaMP7s-WPRE (Addgene, plasmid #104491) into the right VTA. The coordinates for VTA infusions, from bregma, were as follows: anterior-posterior (AP): −3.0 mm; medial-lateral (ML): −0.4 mm; and dorsal-ventral (DV): −4.6 mm. To measure calcium activity in VTA GABA neuronal terminals projecting to the VP, a mono fiber-optic cannula (B280-4604-5.3; Doric Lenses, Canada) was implanted above the right VP (AP: +0.2; ML: -1.2; DV: –4.6). To detect DA release from VTA DAergic neurons, a genetically-encoded fluorescent DA sensor (AAV-hsyn-DA4.4, WZ biosciences) was infused into the NAc shell (AP: +1.2; ML: -0.55; DV: –4.5) of GAD65-Cre mice to ensure that there were no effects of genetic background on photometry measurements, and an optic fiber (B280-4604-5.3; Doric Lenses, Canada) was implanted 0.1 mm above the site of viral infusion.

For optogenetic stimulation experiments, GAD65-Cre mice were infused with AAV2-EF1a-DIO-hChR2(H134R)-eYFP (UNC Vector Core) bilaterally into the VTA (AP: -3.0; ML: ±0.4; DV: -4.6), as previously described (Taylor et al., 2014). Customized bilateral cannulae (B284-2093-5; Doric Lenses, Canada) were implanted in VP (AP: +0.2; ML: ±1.2; DV: -4.6). Before behavioral testing, mice were allowed to recover for at least 28 days.

### Fiber photometry

Methods were adapted from (Crouse, Kim et al. 2020) to measure in vivo calcium and/or dopamine signals as fractional changes of fluorescence (dF/F), using a standard 2-lenses-1-site 405/465 nm fiber photometry system. A console controlled two connectorized LEDs (CLEDs): 405 nm at 208 Hz and 465 nm at 572 Hz, each linked to a five-port Fluorescence MiniCube (FMC5_AE()_AF()_E1()_F1()_S, Doric Lenses). The F1 (500–550 nm) MiniCube port connected to a photoreceiver (AC low mode, New Focus 2151 Visible Femtowatt Photoreceiver, New Focus, CA) via a fiber optic adapter, which linked back to the fiber photometry console. A pigtailed fiber optic rotary joint connected the MiniCube to the patch cord, enabling light transmission in a freely moving mouse through an implanted optical fiber. Optic signals were recorded with Doric Neuroscience Studio (V 5.3.3.14) using Lock-In demodulation at a sampling rate of 12.0 kS/s. Data were decimated by a factor of 100 and saved as a CSV file. For analysis, data were processed with Guided Photometry Analysis in Python (GuPPy)(Sherathiya, Schaid et al. 2021), calculated as a standard z-scored dF/F. The 405 nm fluorescence signal served as a reference to correct artefacts. Baseline fluorescence (F0) was determined by subtracting the fitted 405 nm fluorescence from the 465 nm signal using least linear square regression. The change in fluorescence (dF) was then calculated as the difference between the recording and F0, yielding the final dF/F (%) as a percentage increase. All photometry data are reported as dF/F (%).

### Behavioral Testing

For all behavioral experimental commands and data collection were controlled using Med-PC IV software (command codes available upon request). Except where specified in the text, all animals were food-restricted throughout behavioral sessions. Each mouse was provided with 2 g of chow per day to maintain body weight at 85–90% of its baseline level. Mice were first trained to nose-poke on an FR1 schedule to earn rewards for at least 5 days. A nose-poke in an LED-lit aperture triggered reward delivery (Ensure® Plus, Abbott; ∼24 μl delivered over 2 s) through a reward spout. Mice that earned at least 30 rewards in the last session were randomized and underwent surgery. For Pavlovian conditioning in the free-moving paradigm, a 2-s tone paired with a simultaneous blinking 2-s LED over the receptacle served as the conditioned stimulus (CS). The cue was presented on an RT-60 schedule, and 3 s after each cue ended, a 2-s reward (Ensure) or water was delivered into the receptacle.

Receptacle entries were detected via infrared beam breaks. Each session consisted of 25 reward deliveries. For Pavlovian conditioning experiments, mice were lightly restrained in a custom-made plastic tube with an opening at the end for reward delivery via a metal spout. This setup allowed precise control over the timing of reward delivery and licking/ consumption behavior. A 2-s tone paired with simultaneous 2-s LED blinking above the spout served as the CS. Three seconds after each cue ended, a 2-s reward was pumped into the spout. Each session consisted of 25 reward deliveries. A lickometer recorded the number and timing of licks on the reward spout. Prior to Pavlovian conditioning, mice were acclimated to the restraint paradigm for at least 5 days, 30 min per day. To estimate the minimum number of animals required for statistical conclusions based on the pilot dataset, we performed a paired t-test power analysis using the online calculator at https://www.medcalc.org/en/calc/sample-size-paired-samples-t-test.php.

To assess choice preference for different reward delivery schedules based on reward value, we designed a 60-min PRT, modified from (Milienne-Petiot 2017). The same two juxtaposed nose-poke apertures served as distinct choice options for different reward probabilities. To provide a high reward option, a nose-poke in aperture A led to a 75% chance of receiving a high-value reward (Ensure) and a 25% chance of receiving water (Option A). Conversely, a low reward option was induced with a nose-poke in aperture B, resulting in a 25% chance of Ensure delivery and a 75% chance of water (Option B). Both Ensure and water were delivered via the same composite metal spout. Mice were trained to lick the spout to retrieve rewards. A lickometer recorded the number and timing of licks on the reward spout. To measure preference for each of the two reward schedules, we calculated the ratio of choices for Option B over the total number of effective nose-pokes (that triggered reward delivery), to track positive changes when the preference between the two reward options shifted.

To manipulate the activity of VTA-to-VP GABAergic projecting neurons, a 5-s train of blue light pulses (10 ms, 20 Hz, 465 nm; on-site end power of 90 to 100 mW/mm²) was simultaneously delivered to both sides of the VP through bilateral light cannulae for optogenetic stimulation. Stimulation was triggered when the animal made its first lick on the spout after either a reward or water was delivered following a nose-poke in aperture B.

## Acknowledgments

We thank Dr. Xiao-Bing Gao for intellectual discussions. This study was supported by grants DA014241 and DA036151 from the National Institutes of Health. This work was funded in part by the State of Connecticut, Department of Mental Health and Addiction Services, but this publication does not express the views of the Department of Mental Health and Addiction Services or the State of Connecticut.

