## Supplemental Figures 1-4 for "A ventral tegmental area GABAergic projection to the ventral pallidum regulates value-based decision making in mice"

**This PDF file includes:**

Figures S1 to S4 and their Legends

**Figures**

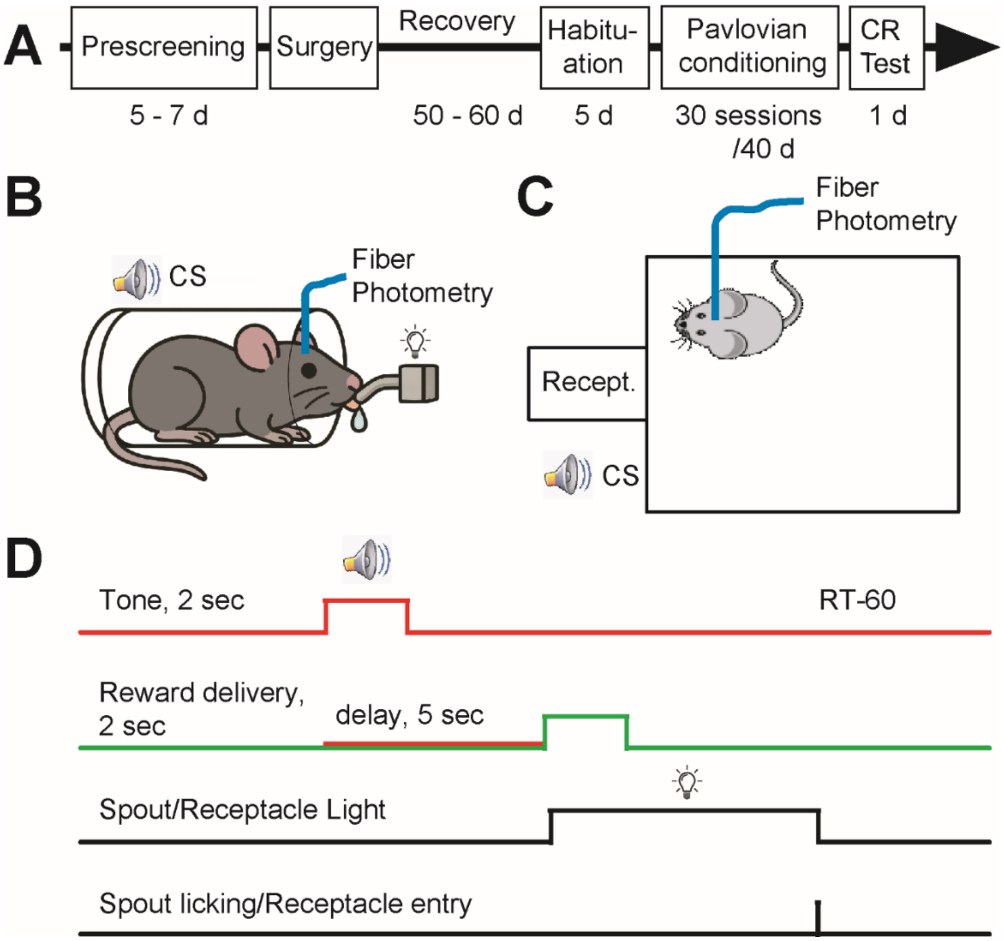


**Fig. S1.** Timeline and paradigms of Pavlovian conditioning procedures. (**A**) Timeline of behavioral procedures used in Pavlovian conditioning. (**B**) Schematic of the body-restraint Pavlovian conditioning paradigm. Mice were placed in a plastic tube to restrain head movement. Upon delivery of a liquid reward to the spout, mice could immediately consume it by licking. If they failed to lick, the reward dropped so the size of the subsequent reward would not be increased. For fiber photometry recording, mice were tethered to a patch cord connected to the fiber photometry system. (**C**) Schematic of the setup used for Pavlovian conditioning in freely-moving mice. Mice were placed in modular test chambers equipped with a wall-mounted receptacle. An auditory tone was generated with a speaker placed outside the chamber. (**D**) Schedule of the Pavlovian conditioning task. Two-second Auditory tones were presented on a RT-60 schedule. A liquid reward was delivered 5 s after tone onset for 2 s. An LED positioned above the spout/ receptacle was illuminated at the time of reward delivery and turned off upon the mouse's first lick following reward onset.

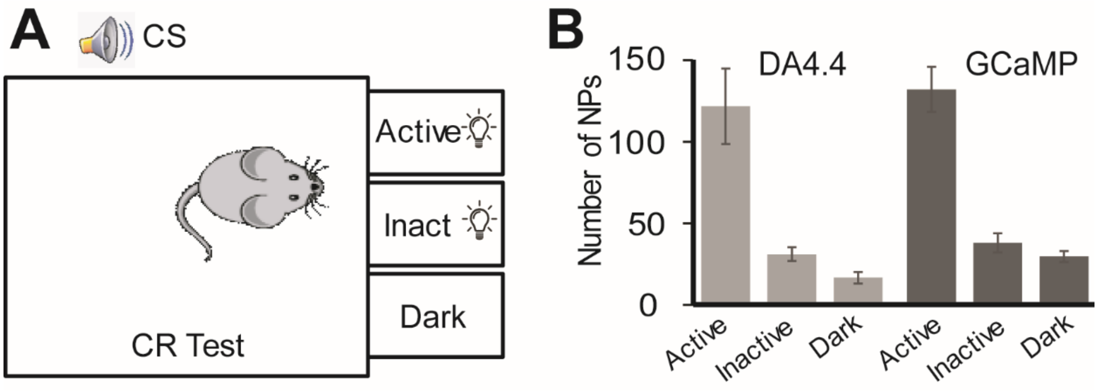


**Fig. S2.** Conditioned reinforcement (CR) test following Pavlovian conditioning. (**A**) Schematic of the behavioral chamber setting used for CR testing, conducted after 25 sessions of Pavlovian conditioning in DA4.4-expressing mice or 30 sessions in GCaMP-expressing mice.A nose-poke into “Active” aperture triggered presentation of the same auditory tone (CS) previously used during Pavlovian conditioning. Nose-pokes into the inactive (Inact) or dark apertures produced no programmed outcome. (**B**) Number of nose-pokes into each aperture during CR testing. The left 3 bars show data from the 4 mice expressing DA4.4 in the NAc shell, tested after 25 conditioning sessions. The right 3 bars show data from the 5 mice expressing GCaMP in VTA GABA neurons, after 30 conditioning sessions. Two-way ANOVA revealed no significant group effect ($F(1,23)=1.36$, $P=0.255$), but a significant main effect (discrimination) between apertures ($F(2,23)=58.1$, $p=1.0\times10^{-9}$), indicating the successful CS-US associative learning during Pavlovian conditioning.

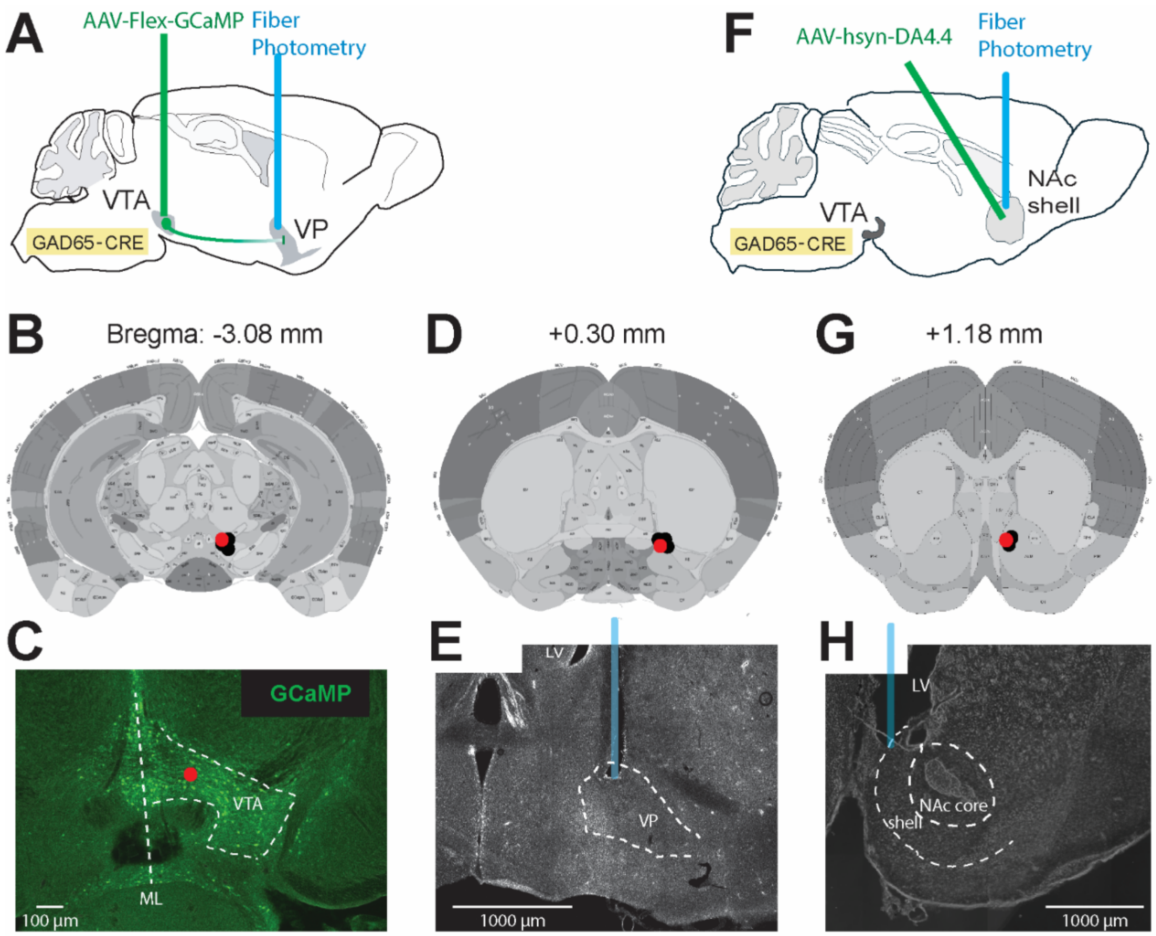


**Fig. S3.** Surgical procedures and verification of viral infusion and fiber placements for fiber photometry. (**A**) Schematic of GCaMP-carrying virus infusion into the VTA and optical fiber implantation targeting the posterior VP to monitor calcium activity via fiber photometry. (**B**)  Histological reconstruction of GCaMP-virus injection sites in the VTA. The red dot corresponds to the injection site shown in panel C. (**C**) A representative confocal image showing GCaMP expression in the VTA following behavioral experiments. (**D**) Fiber tip placements in the VP for photometry recordings. Red and black dots indicate individual fiber tip locations across subjects. The red dot corresponds to the fiber placement shown in panel E. (**E**) A representative image showing the actual placement of an optical fiber with the tip right above the posterior VP, corresponding to the red dot in penal D. (**F**) Schematic of DA4.4-carrying virus infusion into the NAc shell and optical fiber implantation for photometric monitoring of DA transmission, with the tip positioned right above the virus injection site. (**G**) Histological reconstruction of DA4.4-virus injection sites in the NAc shell. The red dot corresponds to the fiber placement shown in panel H. (**H**) A representative image showing the actual placement of the optical fiber tip above the viral injection site in the NAc shell, corresponding to the red dot in panel D.

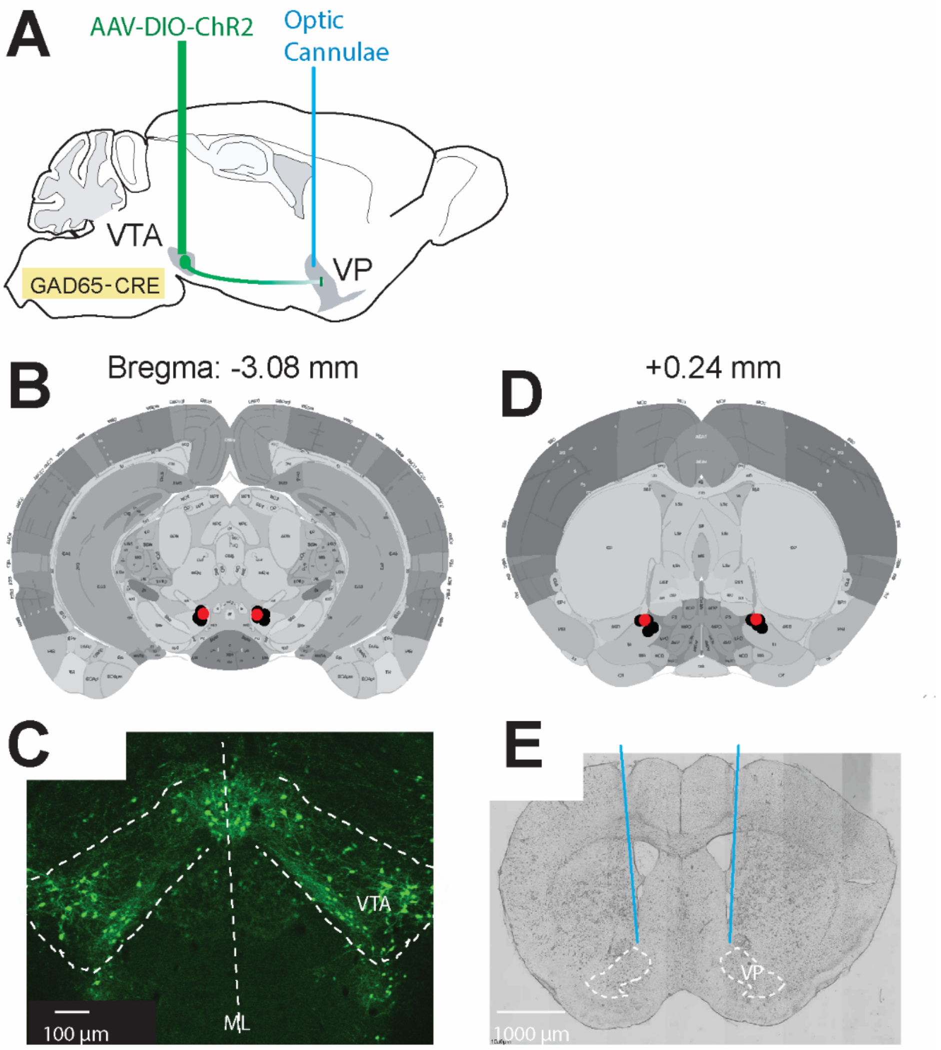


**Fig. S4.** Surgical procedures and verification of viral infusion and optical cannulae placements for optogenetic stimulation. (**A**) Schematic of ChR2-carrying virus infusion into the VTA and optical cannulae implantation targeting the posterior VP for stimulating VTA-to-VP projecting axons terminals. (**B**) Histological reconstruction of ChR2-virus injection sites in the VTA. (**C**) A representative confocal image showing ChR2 expression in the VTA after completion of behavioral experiments. (**D**) Schematic showing optical cannula tip placements in the VP. Red and black dots indicate individual cannula tip locations. (**E**) A representative image showing the actual placement of an optical fiber with the tip positioned right above the posterior VP, corresponding to the red dot in penal D.
